# Metabolic Targeting of SREBP1 Reprograms the Obesity-Driven Ascites Immune Microenvironment and Enhances Chemotherapy Response in Obesity-Associated Ovarian Cancer

**DOI:** 10.64898/2026.09.14.751517

**Authors:** Jing Yang, Tyvette S. Hilliard, Yueying Liu, Emily Richardson, Sophia Santoso, Viktoriia Khomenko, Molly Conroy, Brandon Martin, Emily Cronberger, Gena Dominique, Zhikun Wang, Jeffrey Johnson, Wanrui Wang, Ijeoma Asilebo, M. Sharon Stack

## Abstract

Obesity has an adverse effect on survival of women with ovarian cancer (OC), supporting a link between obesity, metastatic progression, and therapeutic response, but comprehensive mechanistic insight is lacking. In pre-clinical models of diet-induced obesity and OC, mice fed a high fat diet (HFD) exhibit substantially increased tumor burden relative to low fat diet (LFD) mice and respond poorly to standard-of-care (SOC) chemotherapy. Both HFD murine tumors and tumors from women with BMI>30 show increased expression and nuclear localization of sterol regulatory element binding protein 1 (SREBP1), a master regulator of lipogenesis and lipid transport. The objective of this study was to assess whether inhibition of SREBP1 processing could improve SOC chemotherapy response in obesity-associated OC. We evaluated the FDA-approved drug Nelfinavir (NFV), a pharmacologic inhibitor of site-2 protease that blocks SREBP1 processing, in combination with SOC (paclitaxel+carboplatin) chemotherapy in preclinical models of diet-induced obesity and OC. Our results demonstrate significantly improved therapeutic response, corresponding changes in intra-tumoral adipocytes, and reprogramming of the obesity-driven ascites immune microenvironment. These findings support a model in which NFV-mediated inhibition of SREBP1 processing disrupts obesity-associated metabolic pathways linked to immune suppression and therapeutic resistance, thereby enhancing response to SOC chemotherapy in obesity-associated OC. Collectively, these results highlight metabolic targeting as a strategy to improve therapeutic outcomes in obesity-associated ovarian cancer.

## INTRODUCTION

Ovarian cancer (OC) is the most lethal and second most prevalent gynecologic malignancy in the United States (1). In 2025, OC is anticipated to result in approximately 12,730 deaths in the United States and 207,000 deaths worldwide (1,2). The high mortality rate is primarily due to late-stage diagnosis (FIGO stages III and IV), extensive intraperitoneal metastasis, and significant resistance to chemotherapy. The majority of women with epithelial ovarian cancer are diagnosed with metastatic disease, resulting in a poor 5-year survival of ∼49% across all stages and ∼31% for those diagnosed with distant metastatic disease (3). Clearly, novel approaches are needed to provide a comprehensive understanding of tumor-host interactions and identify novel targets for improved survival of women with OC.

OC metastasis is initiated via direct extension of the primary fallopian tube or ovarian tumors, and exfoliation of cells and multicellular aggregates into the peritoneal cavity where dissemination is facilitated by ascites fluid (4–6). The presence of large volumes of malignant ascitic fluid provides a unique tumor microenvironment that promotes progression and metastasis, enables immune evasion, and mediates therapy resistance (4–8). Ovarian tumors are genetically highly heterogeneous and exhibit diffuse patterns of intraperitoneal dissemination with preferential metastasis to the adipose-rich omental fat pad and multiple sites within the peritoneal cavity (6).

Compounding a poor prognosis is obesity, a recognized non-infectious pandemic that affects >40% of adult women in the United States (9). We have previously demonstrated a positive correlation between obesity and OC metastatic success in multiple murine pre-clinical models of diet induced obesity (DIO) and mutational obesity (*Ob/Ob* mice) (10). This was characterized by significantly enhanced intraperitoneal metastatic tumor burden and an altered immune microenvironment reflected by a decreased M1/M2 macrophage ratio in obese hosts (10). Moreover, relative to mice fed a low fat diet (10% fat), mice on a high fat diet (45% fat) demonstrated a poor response to standard-of-care (SOC) chemotherapy (paclitaxel and carboplatin, PC), characterized by substantial residual tumor burden at the end of the treatment cycle (11). This is consistent with meta-analyses that show a relationship between obesity and OC survival in women with tumors of serous, endometrioid and mucinous histology, implicating a link between host obesity, metastatic success, and chemotherapy response (12–22).

Our published data also demonstrate that DIO resulted in elevated expression and nuclear localization of sterol regulatory element binding protein 1 (SREBP1) (10). SREBP1, a transcription factor and master regulator of *de novo* lipogenesis, promotes cancer cell proliferation and metastasis through lipogenic reprogramming (6,23). SREBP1 is present in the endoplasmic reticulum (ER) as an inactive precursor that undergoes sequential regulated intramembrane proteolysis, catalyzed by site-1 protease and site-2 protease in the Golgi, to release a mature form designated nSREBP1. This form complexes with importin β, is transported to the nucleus as a dimer, and binds to sterol regulatory element sequences in the promoters of target genes involved in synthesis and uptake of cholesterol, fatty acids, triglycerides, and phospholipids. Thus SREBP1 can enhance cancer growth and metastasis via lipogenic reprogramming, allowing the metabolic demands of cancer to be met by serving as fuel sources and providing precursors for cell membrane synthesis for rapidly replicating cancer cells (6,23).

Emerging evidence further suggests that SREBP1-mediated metabolic reprogramming contributes to the establishment of an immunosuppressive tumor microenvironment. In both tumor and immune cells, dysregulated lipid metabolism has been linked to impaired anti-tumor immunity, altered macrophage polarization, and therapeutic resistance (23–26). These observations suggest that elevated SREBP1 activity may represent a mechanistic link between obesity-associated metabolic dysfunction, immune suppression and poor response to chemotherapy in OC. Since transcription factors are challenging to target, several compounds have been identified to indirectly block SREBP1 activity by inhibiting proteolytic processing and activation. Among these is Nelfinavir (NFV), an FDA-approved HIV protease inhibitor, that has been shown to inhibit SREBP1 processing and downstream lipogenic signaling while exhibiting anti-tumor activity in multiple cancer models (27–29). Therefore, repurposing NFV in combination with SOC chemotherapy represents a rational approach to simultaneously target metabolic pathways associated with tumor progression, immune suppression and treatment resistance.

In this study, we have evaluated the effect of two pharmacologic inhibitors of SREBP1 processing, Fatostatin and NFV, in combination with SOC chemotherapy (paclitaxel and carboplatin) in pre-clinical models of OC and DIO. Our results show significantly improved therapeutic response and diminished post-treatment tumor recurrence in mice receiving combination therapy with SOC and NFV. Corresponding changes in intra-tumoral adiposity and in the obesity-driven ascites immune landscape provide mechanistic insight into how NFV-mediated inhibition of SREBP1 signaling enhances chemotherapy efficacy and improves therapeutic outcomes in obesity-associated ovarian cancer.

## MATERIALS AND METHODS

### Immunohistochemical analysis of human ovarian tumors

De-identified tumor blocks from women diagnosed with high-grade serous ovarian cancer, with a defined body mass index (BMI), were purchased from the Indiana University Simon Cancer Center tissue bank (Indianapolis, IN). Tissues were accrued with informed consent in accordance with relevant guidelines and regulations of the Indiana University School of Medicine Institutional Review Board. This study using de-identified tissue was conducted with the approval of the University of Notre Dame Institutional Review Board in accordance with relevant guidelines and regulations. A summary of patient data is provided in **Suppl. Table 1**.

Tumor sections were subjected to immunohistochemical analysis for SREBP-1 (Abcam, Cat# ab191857, 1:500 dilution, Waltham, MA), or ‘no primary antibody’ controls, as previously described (10). The detection was facilitated by a peroxidase-conjugated secondary antibody (Vector Laboratories, Cat# MP-7401, Newark, CA) and a DAB peroxidase substrate (Vector Laboratories, Cat# SK-4105). Slides were digitized using the Aperio ScanScope CS (Leica Biosystems Inc., Buffalo Grove, IL) and subsequently uploaded to the eSlide Manager Database. Nuclear SREBP-1 staining was quantified by analyzing a minimum of 400 epithelial tumor cells per slide by an investigator blinded to BMI status (M.S.S.). Statistical analyses were performed using Student’s t-test, with a p-value < 0.05 considered statistically significant. All analyses were conducted in GraphPad Prism software (version 10.6.1; GraphPad Software, San Diego, CA; www.graphpad.com).

### Pre-clinical therapeutic trials in an ovarian cancer allograft model of mice with diet-induced obesity

All pre-clinical murine studies were conducted with approval of the University of Notre Dame’s Institutional Animal Care and Use Committee (IACUC) in accordance with relevant guidelines and regulations. Female C57Bl/6 mice (3-6 weeks old, The Jackson Laboratory, Bar Harbor, ME) were maintained on a high-fat diet (HFD; 45% fat, Research Diets, Cat# D12451, New Brunswick, NJ) until they reached a weight of over 30g (15-30 weeks). The mice continued on this high-fat diet for the duration of the study.

Allograft studies utilized the murine cell line ID8*Trp53*^-/-^, which is syngeneic to C57Bl/6 mice, and harbors a *Trp53* deletion (30). These cells were generously provided by Dr. Ian McNeish (Glasgow, UK) and were transduced to express red fluorescent protein (RFP) as previously described (31). ID8*Trp53*^-/-^-RFP cells were cultured in Dulbecco’s Modified Eagle Medium (Corning, Cat# 10-014-CV, Corning, NY) containing 4% Fetal Bovine Serum (Thermo Fisher Scientific, Cat# A5256701, Grand Island, NY), 1% Penicillin/Streptomycin (Thermo Fisher Scientific, Cat# 15140122), and 1% Insulin-transferrin-sodium selenite (Sigma-Aldrich, Cat# I1884, St. Louis, MO).

Mice were injected intra-peritoneally (i.p.) with ID8*Trp53*^-/-^-RFP cells (1×10^6^ in 1 ml PBS). Following injection, mice were imaged weekly under isoflurane anesthesia using the IVIS Lumina *in vivo* imaging system. After approximately 3 weeks, when sufficient tumor burden had developed, mice were randomized into two cohorts: “standard-of-care (SOC)” and “combination therapy.” The SOC cohort received weight-adjusted chemotherapy consisting of paclitaxel (6 mg/kg, Sigma-Aldrich, Cat# T7191) and carboplatin (15 mg/kg, Sigma-Aldrich, Cat# 1096407) for 6 cycles (twice weekly i.p. for 3 weeks),designated as PC. The combination therapy cohort received the same SOC chemotherapy along with an additional SREBP-1 targeting agent. Treatment compounds included Nelfinavir (Fisher Scientific, Cat# N0986100MG, Pittsburgh, PA) at 50 mg/kg for 9 cycles (thrice weekly i.p., 3 weeks), Nelfinavir at 100 mg/kg for 18 cycles (thrice weekly i.p., 6 weeks), or Fatostatin (Sigma-Aldrich, Cat# 341329) at 30 mg/kg for 9 cycles (thrice weekly i.p., 3 weeks).

Throughout the study, mice were closely monitored for signs of weight loss, lethargy, and ascites accumulation. Mice were euthanized for endpoint dissection one week following the last treatment using the Euthanex system for CO2 inhalation followed by cervical dislocation in accordance with University of Notre Dame IACUC guidelines and regulations. Ascites was immediately collected using a syringe. The peritoneal cavity was accessed through incisions made along the midline and sides of the ventral parietal peritoneum. Abdominal tumor burden was assessed by scanning the abdominal organs *in situ* using the IVIS Lumina *in vivo* imaging system, followed by the removal of individual organs for *ex vivo* imaging (10,11,31,32). Total abdominal or organ-specific tumor burden was quantified using ImageJ by calculating the tumor area and the intensity of the RFP signal (Raw Integrated Density).

Statistical analyses were performed using Student’s t-test, with a p-value < 0.05 considered statistically significant. All analyses were conducted in GraphPad Prism. All animal procedures were performed in accordance with the regulations and approval of the Institutional Animal Care and Use Committee at the University of Notre Dame.

### Histology and immunohistochemical analysis of murine ovarian tumors

Abdominal organs were fixed in 10% neutral-buffered formalin (Fisher Scientific, Cat# 22-050-105) and subsequently paraffin-embedded for histological analysis. Tissue sections were stained with hematoxylin (Fisher Scientific, Cat# 22-050-206) and eosin (Fisher Scientific, Cat# SE23-500D) according to standard H&E protocols (10,11).

For immunohistochemistry, sections were deparaffinized and endogenous peroxidase activity was quenched with 3% hydrogen peroxide for 30 minutes. Antigen retrieval was performed by boiling in 10 mM sodium citrate (pH 6.0) for 30 minutes. To reduce nonspecific binding, sections were blocked with 2.5% normal horse serum in PBS for 1 hour at room temperature. Slides were then incubated overnight at 4°C with primary antibodies diluted in 2.5% normal horse serum. Control slides omitted the primary antibody. The following antibodies were used: PCNA (Cell Signaling Technology, Cat# 13110S,1:12,000 dilution; Danvers, MA); SREBP-1 (Abcam, Cat# ab191857, 1:500 dilution); Phospho-Histone H2A.X (Cell Signaling Technology, Cat# 9718S, 1:480 dilution); iNOS (Abcam, Cat# ab15323, 1:1,000 dilution); CD206 (Abcam, Cat# ab64693, 1:6,400 dilution). The detection was facilitated by a peroxidase-conjugated secondary antibody (Vector Laboratories, Cat# MP-7401) and a DAB peroxidase substrate (Vector Laboratories, Cat# SK-4105).

Slides were digitized using the Aperio ScanScope CS system (Leica Biosystems) and archived in the eSlide Manager database. Tumor and adipose tissue regions were annotated and analyzed with the percent-positive macro algorithm in Aperio ePathology ImageScope to quantify DAB chromogen-positive (brown) cells. Statistical analyses were performed using Student’s t-test or the Mann-Whitney test, with p<0.05 considered statistically significant. All analyses were conducted in GraphPad Prism (version 10.6.1).

### Isolation and multiplex flow cytometric analysis of murine immune cells

Peritoneal immune cells were collected by sacrificing mice and injecting 6 mL of PBS into the peritoneal cavity. After agitation for ∼1 minute, all fluid (including ascites and lavage) was recovered and centrifuged at 450g for 5 minutes at 4 °C. For spleen cell preparation, spleens were mechanically dissociated in 2 mL DMEM using a 40 µm cell strainer placed in a petri dish until no dark red tissue remained. Cells were collected and centrifuged at 300g for 3 min at 4 °C.

Cell pellets (from peritoneal lavage or spleen) were resuspended in ACK lysis buffer (Gibco, Cat# A1049201, Grand Island, NY) to remove red blood cells, followed by neutralization with cold PBS. After washing, cells were resuspended at 1×10^6^ cells/mL in PBS and stained with the LIVE/DEAD Fixable Yellow Dead Cell Stain Kit (Thermo Fisher Scientific, Cat# L34959, Eugene, OR; 0.25 µL/mL) for 30 minutes in the dark. Cells were then washed, resuspended in PBS containing 10% FBS, and plated at 1×10^6^ cells/well in 96-well V-bottom plates. To block nonspecific Fc receptor binding, samples were incubated with anti-CD16/CD32 antibody (BioLegend, Cat# 156604, San Diego, CA) for 15 minutes in the dark. Cells were subsequently stained with a cocktail of fluorescently conjugated antibodies for 15 minutes protected from light. After three washes, cells were resuspended in 200 µL PBS containing 10% FBS and analyzed using a Cytek Northern Lights flow cytometer (Cytek Biosciences, NL-2000, Bethesda, MD) equipped with blue and violet lasers and SpectroFlo software. Data were acquired at a medium flow rate with a threshold of 50,000 P1-gated events, using live spectral unmixing against single-stained peritoneal lavage controls.

Flow cytometry data were analyzed with FlowJo software (version 11.1; BD Biosciences, Ashland, OR). Gating was performed on single live cells, and populations were quantified as a percentage of total live parental cells. Statistical analyses were performed using Student’s t-test or the Mann-Whitney test, with p<0.05 considered statistically significant. All analyses were conducted in GraphPad Prism (version 10.6.1; GraphPad Software). Antibodies used are listed in **Suppl. Table 2.**

## RESULTS

### SREBP1 expression in human ovarian tumors correlates with body mass index (BMI)

We have previously observed a striking upregulation of nuclear-localized SREBP1 staining in tumors from a DIO study in which mice were fed a western diet (40% fat), relative to control diet (10% fat) mice (10). Similar results were observed in tumors grown in *Ob/Ob* mice relative to wild-type controls (10). SREBP1 staining showed intense nuclear localization, indicating transcriptional activity that contributes to lipogenic reprogramming of tumor cells. To evaluate whether human tumors exhibit similar patterns of SREBP1 expression, high grade serous ovarian tumors from women with a range of BMIs were examined [**Suppl. Table 1**]. Immunohistochemical analysis shows that tumors from obese women (BMI>30, n=10) exhibit a similar pattern of strong nuclear SREBP1 staining relative to tumors from women with BMI<30 (n=10) [**Fig. 1A-E**]. A significant difference in SREBP1-positive nuclei was observed between the low BMI and high BMI groups [**Fig. 1G**, p<0.001] and SREBP1-stained nuclei positively correlated with BMI [**Fig. 1H**, r^2^=0.72].

**Fig. 1.**
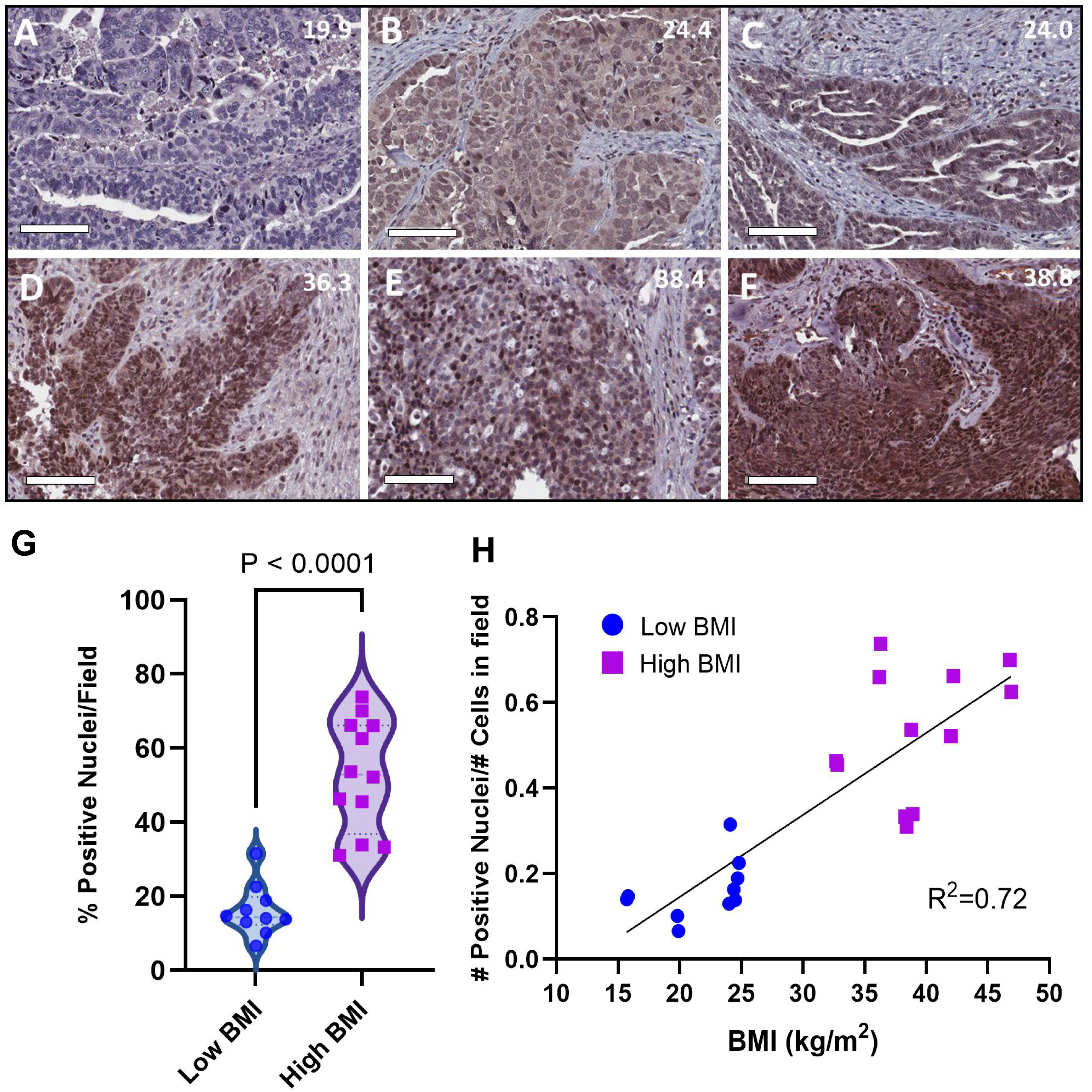
SREBP1 nuclear localization is enhanced in OC tumors from women with high body mass index (BMI). **(A-F)** Tumors (n=20) from women with low BMI and high BMI were subjected to immunohistochemical staining for SREB1 as described in Methods. Representative images are shown, with the patient’s BMI displayed in the upper right corner of each panel. Scale bar: 100 μm. **(G)** Comparison of nuclear SREBP1 expression in ovarian tumors from patients with low BMI (<30) *vs* high BMI (>30). **(H)** Relationship of nuclear SREBP1 expression with BMI in ovarian cancer patients.

### Combination treatment with SOC chemotherapy and inhibitors of SREBP1 processing in the DIO setting

Human OC tumors from women with high BMI and tumors from OC-bearing mice on a high fat diet exhibit enhanced nuclear SREBP1 staining (10). Moreover our pre-clinical murine studies show poor response to standard-of-care chemotherapy (paclitaxel and carboplatin, PC) in the high fat diet setting relative to low fat diet mice (11); therefore a pilot pre-clinical combination therapy trial was conducted. DIO was generated by feeding mice a high fat diet (45% fat) until a weight of ∼30g was achieved, followed by i.p. injection of RFP-tagged ID8-*Trp53*^-/-^ cells [**Fig. 2A**]. After three weeks of tumor growth, mice were randomized to one of three treatment groups [**Fig. 2A**, n=6/group]. Control mice received standard-of-care chemotherapy (paclitaxel and carboplatin, designated PC) twice weekly for 3 weeks. Combination therapy cohorts received PC together with an inhibitor of SREBP1 processing, either Fatostatin or NFV. Fatostatin is a diarylthiazole derivative that blocks SREBP1 translocation from the ER to the Golgi, thus inhibiting proteolytic processing and maturation of SREBP1 (6). NFV (Viracept) is an inhibitor of site-2 protease activity, resulting in suppressed intramembrane proteolysis of SREBP1, thus blocking nuclear translocation (27,28). Mice in the PC+Fatostatin cohort showed severe dose-limiting toxicity characterized by extremely rapid and substantial weight loss [**Suppl. Fig. 1A**], hair loss, and severe dermatitis (*not shown*), and this arm of the study was thus discontinued at week 6. In contrast, PC+NFV was well-tolerated, as the weight of subjects in this arm did not differ significantly from the PC alone arm. The study was repeated with only the PC and PC+NFV arms (n=12/group). Following three weeks of treatment (weeks 3-6), mice in the PC+NFV group showed a similar initial treatment response when compared to mice receiving PC alone [**Fig. 2B,D**]. However, post-treatment recurrence at week 9 was significantly diminished in the PC+NFV cohort relative to PC controls [**Fig. 2C,E**], suggesting that inhibition of SREBP1 processing delays recurrence in the DIO setting.

**Fig. 2.**
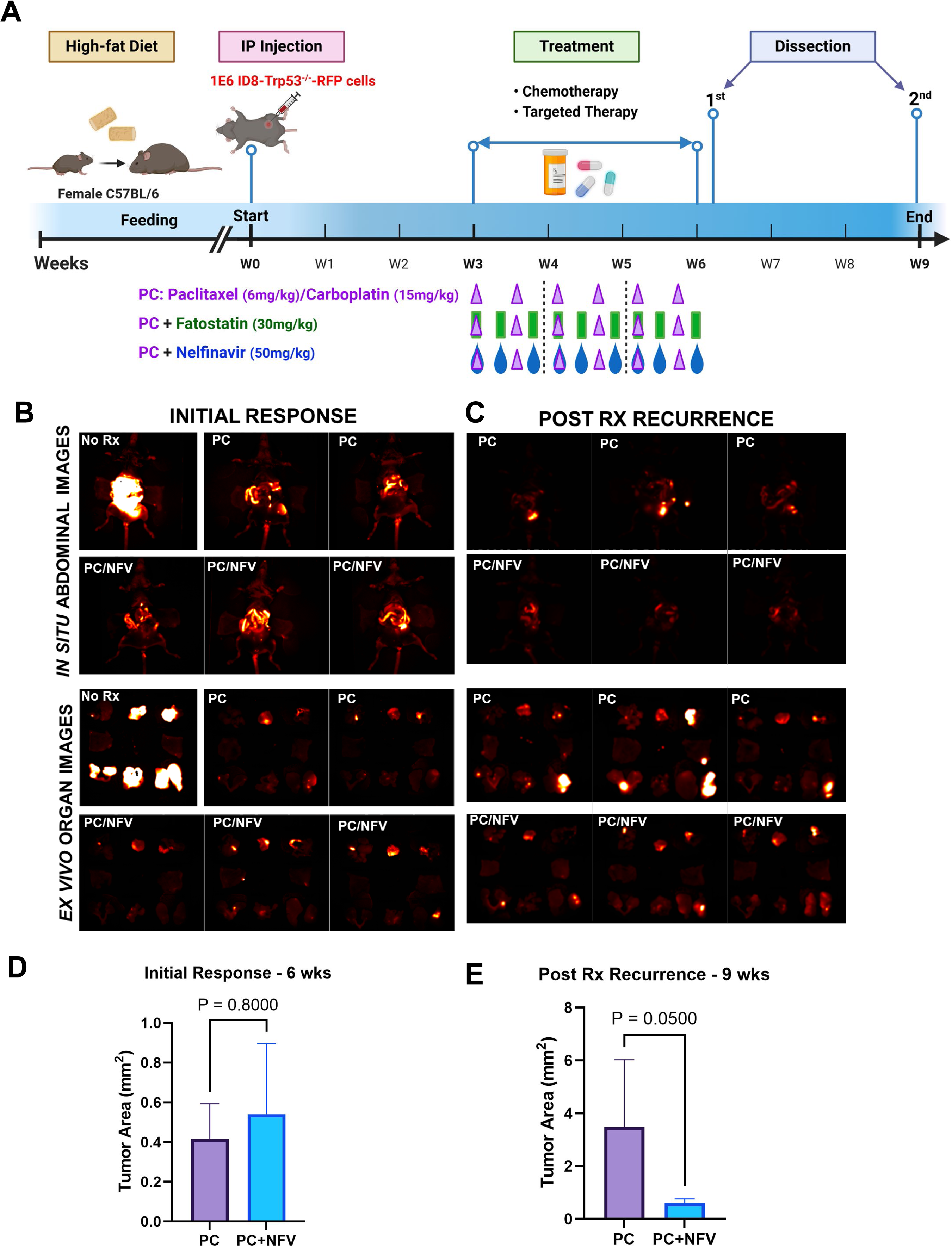
Preliminary murine pre-clinical trial comparing standard-of-care with and without SREBP1-targeting combination therapy. **(A)** Overview of initial study design. All mice (n=6/group) were fed a high-fat diet (HFD, 45% fat) until they reached a weight of 30g, then injected with RFP-tagged ID8-*Trp53*^-/-^ cells (10^6^) to establish tumor burden. Tumor-bearing mice were monitored by longitudinal *in vivo* imaging for development of equivalent tumor burden, then treated either with standard-of-care Paclitaxel (6mg/kg) & Carboplatin (15mg/kg)] (designated PC, twice weekly for 3 weeks); PC + Fatostatin (30mg/kg, thrice weekly for 3 weeks); or PC + Nelfinavir (designated NFV, 50mg/kg, thrice weekly for 3 weeks). The PC+Fatostatin arm was discontinued at week 6 due to severe toxicity (**Suppl.** Fig. 1). **(B-E)** The study was repeated with only the PC and PC+NFV arms. Mice (n=12/group) were dissected following treatment at **(B,D)** week 6 to monitor initial response or at **(C,E)** week 9 to monitor post-treatment recurrence. The upper panels in B and C show *in situ* abdominal tumor burden while the lower panels depict *ex vivo* organ-specific tumor burden. Tumor burden was evaluated by quantitative fluorescence imaging using the Caliper IVIS Lumina II multispectral *in vivo* imaging system and quantified using ImageJ. No Rx = untreated tumor-bearing mice.

Based on the results above, the PC+NFV trial (n=10/group) was repeated with the NFV dose doubled in concentration and cycle number [**Fig. 3A**], with the NFV cohort continuing to receive thrice weekly NFV alone for three additional weeks (for a total of 6 weeks) following the initial six cycles of PC+NFV. Mice were evaluated at week 9 following tumor injection for quantitation of tumor burden. The higher dose of NFV was well-tolerated [**Suppl. Fig. 1B**]. Moreover, mice in the PC+NFV cohort showed a significantly improved treatment response compared to those in the cohort treated with PC alone [**Fig.3B-D**], suggesting that higher dose inhibition of SREBP1 processing improves initial response to standard of care chemotherapy in the DIO setting.

**Fig. 3.**
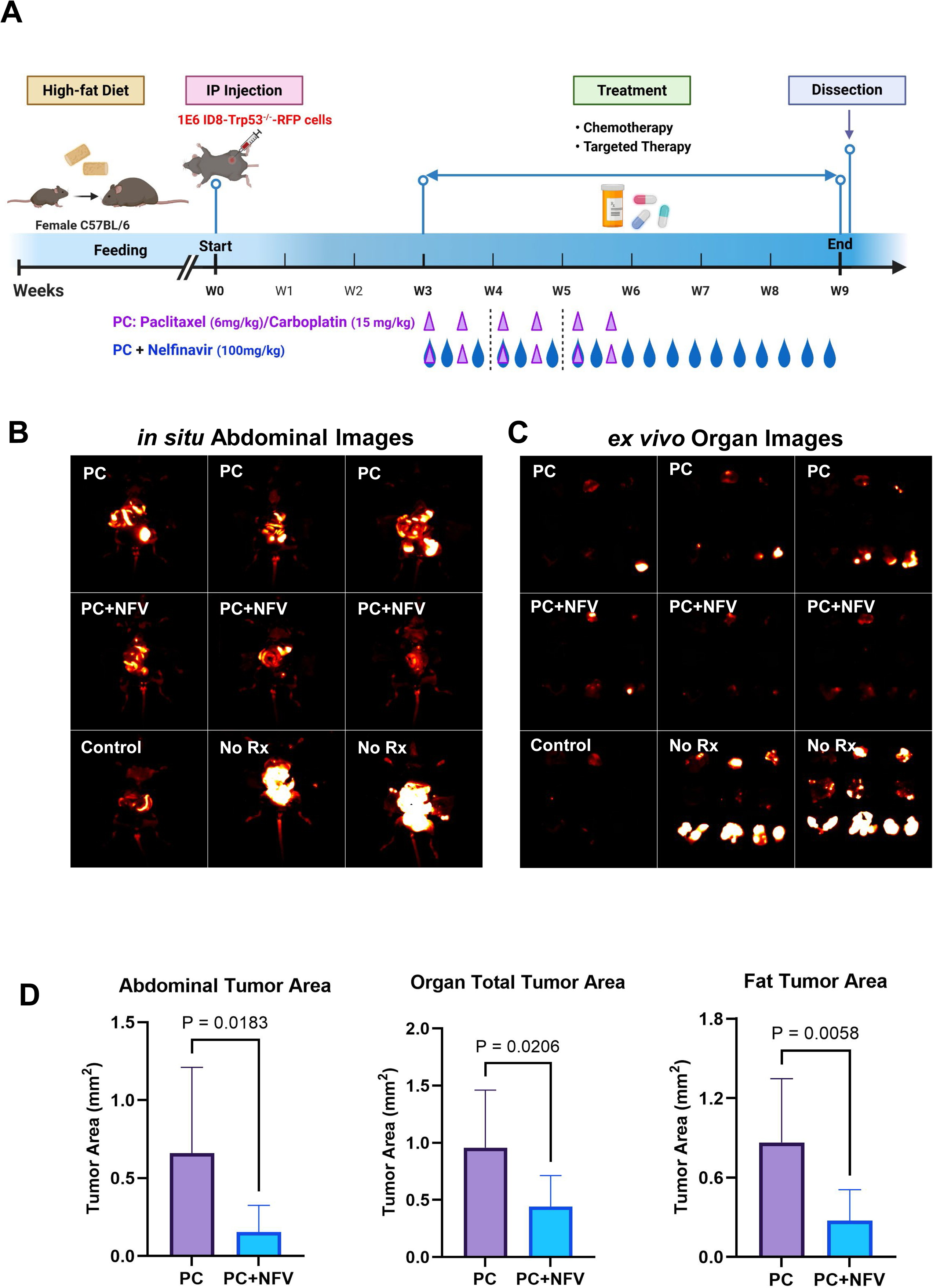
Pre-clinical trial comparing standard-of-care with and without Nelfinavir. (A) Overview of study design. Mice (n=10/group) were fed a high-fat diet (HFD, 45% fat), then injected with RFP-tagged ID8-*Trp53*^-/-^ cells (10^6^) to establish tumor burden. Tumor-bearing mice were monitored by longitudinal *in vivo* imaging for development of equivalent tumor burden, then treated either with standard-of-care Paclitaxel (6mg/kg) & Carboplatin (15mg/kg)] (designated PC, twice weekly for 3 weeks) or PC + Nelfinavir (designated NFV, 100mg/kg, thrice weekly for 6 weeks). **(B-C)** Mice (n=10/group) were dissected following treatment cessation at week 9 to assess therapeutic response. Panel B shows *in situ* abdominal tumor burden and panel C depicts *ex vivo* organ-specific tumor burden. **(D)** Tumor burden was evaluated by quantitative fluorescence imaging using the Caliper IVIS Lumina II multispectral *in vivo* imaging system and quantified using ImageJ. Graphs depict overall abdominal tumor area (*in situ*), sum of individual organs tumor areas (*ex vivo*) and adipose-only tumor area. No Rx = untreated tumor-bearing mice. Control = tumor naïve mice included as an imaging control to account for tissue and organ autofluorescence.

### Immunohistochemical analysis of tumors

Immunohistochemical analysis of tumors from treatment groups shows that subjects treated with PC+NFV exhibited significantly less nuclear-localized SREBP1 in both tumor cells (**Fig. 4A**) and in cancer-associated adipocytes (**Fig. 4B**) relative to mice treated with PC alone, indicating that NFV inhibited SREBP1 processing and the subsequent nuclear translocation *in vivo*. Overall numbers of intra-tumoral adipocytes were decreased in the PC+NFV cohort (**Fig. 4C**) and adipocyte size was also significantly decreased in this group (1078+/-768 mm^2^) relative to those from mice treated with PC only (2193+/-1706 mm^2^, p<0.001). Decreased expression of PCNA in the combination therapy cohort suggests that NFV enhanced the chemotherapy response in the DIO murine model, resulting in a lower rate of cancer cell division and DNA synthesis (**Fig. 4D**) (33). This is supported by decreased expression of the phosphorylated histone protein H2AX in the combination therapy group, indicative of reduced residual DNA damage in this cohort (**Fig. 4E**).

**Figure 4.**
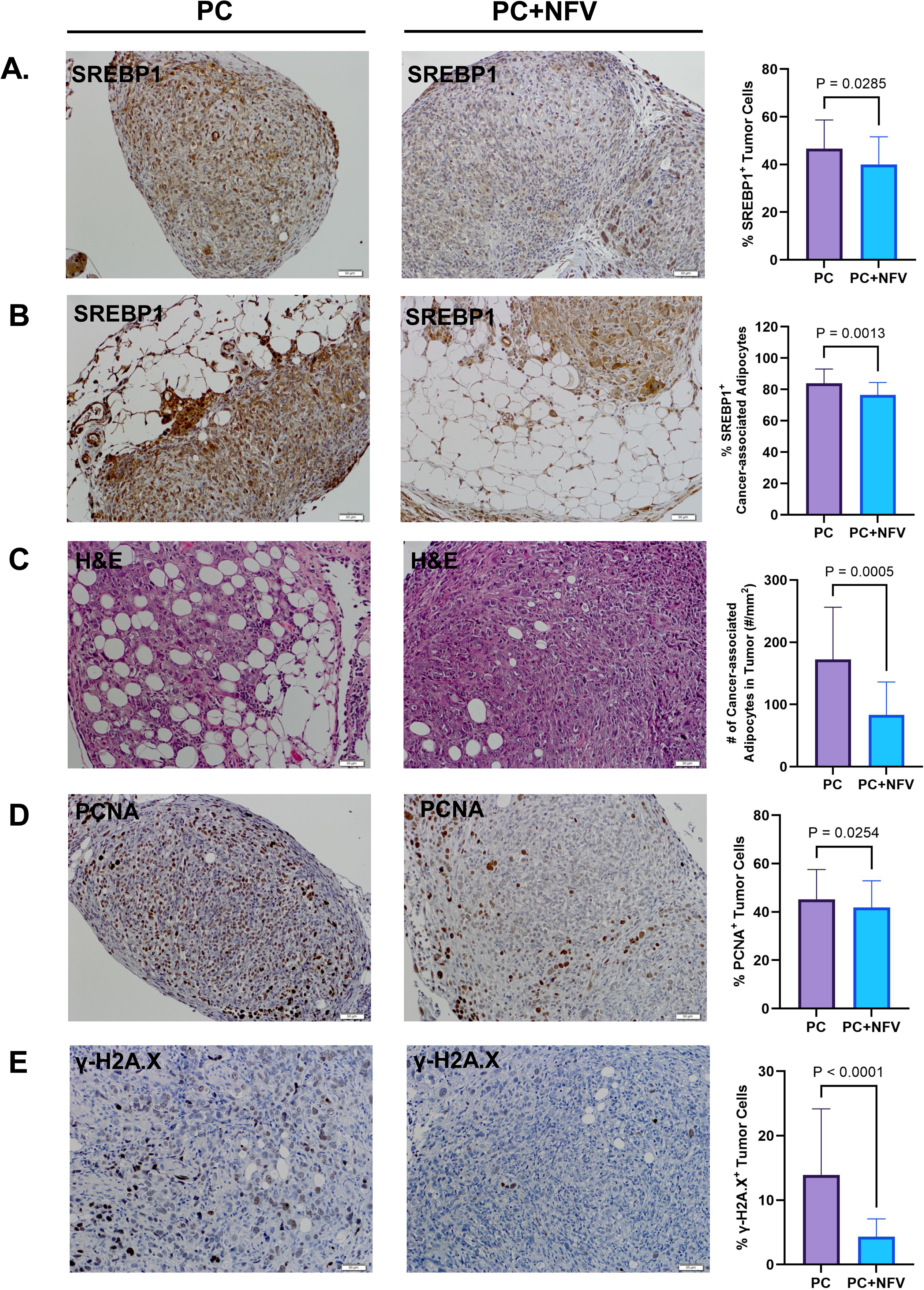
Immunohistochemical analysis of murine tumor tissues. **(A,B)** Tumor sections were subjected to immunohistochemical staining for SREBP1, with nuclear expression in **(A)** tumor cells and **(B)** cancer-associated adipocytes. **(C)** H&E staining of tumor sections highlighting cancer-associated adipocytes. **(D,E)** Immunohistochemical staining of **(D)** PCNA (proliferation) and **(E)** γ-H2AX (DNA damage). Antigen expression was measured using Aperio ImageScope and pairwise statistical comparisons were conducted using Student’s t-test (GraphPad). Scale bar: 50 μm.

### Combination therapy alters the peritoneal immune landscape

In a previous study using immunophenotyping of human tumors from women with a range of body mass indices (BMI, 20-40 kg/m^2^), we reported that tumors from high BMI patients (>30 kg/m^2^) exhibited a decreased M1/M2 macrophage ratio relative to tumors from women with low BMI (<30 kg/m^2^) (11). Similar results were obtained in pre-clinical murine models, as tumors from mice on a high fat diet protocol also demonstrated a significantly decreased M1/M2 relative to mice on a low fat diet (11). In the current study, immunohistochemical analysis of tumors and cancer-associated adipocytes showed no significant changes in iNOS-positive M1 macrophages between high fat diet mice treated with either PC or PC+NFV [**Fig. 5A,B**], while staining for CD206 showed significantly decreased M2 macrophage levels in the PC+NFV cohort. Overall, PC+NFV combination therapy increased the tumor M1/M2 macrophage ratio by 1.5-2.7-fold relative to the PC only cohort. Similar results were observed in peritoneal lavage/ascites using multiplex flow cytometry to obtain peritoneal immune profiles, showing a 2.9-5.0-fold increase in M1/M2 ratios in PC+NFV-treated mice relative to those treated with PC alone [**Fig. 5C**]. Mice that were treated with the PC+NFV combination therapy also had significantly diminished levels of T-regulatory cells (Tregs) [**Fig. 6A**], innate lymphoid cells (ILCs) [**Fig. 6B**] and conventional type 1 dendritic cells (cDC1s) [**Fig. 6C**] and greater numbers of γδ-T cells [**Fig. 6D**], conventional type 2 dendritic cells (cDC2s) [**Fig. 6E**], polymorphonuclear myeloid-derived suppressor cells (PMN-MDSCs) [**Fig. 6F**] and mast cells [**Fig. 6G**]. To distinguish local tumor-associated immune responses from systemic immune responses to the treatment, splenic immune populations were also analyzed. Combination therapy significantly reduced Tregs, natural killer (NK) cells, total dendritic cells, cDC1s, and both monocytic and polymorphonuclear myeloid-derived suppressor cells (Mo-MDSCs and PMN-MDSCs) while increasing mast cells and cDC2s in the spleen. Full ascites and splenic immune profiles are provided in **Suppl. Fig. 2A,B** respectively. These data indicate PC+NFV treatment induces local and systemic immune remodeling across tumors, ascites, and spleen.

**Figure 5.**
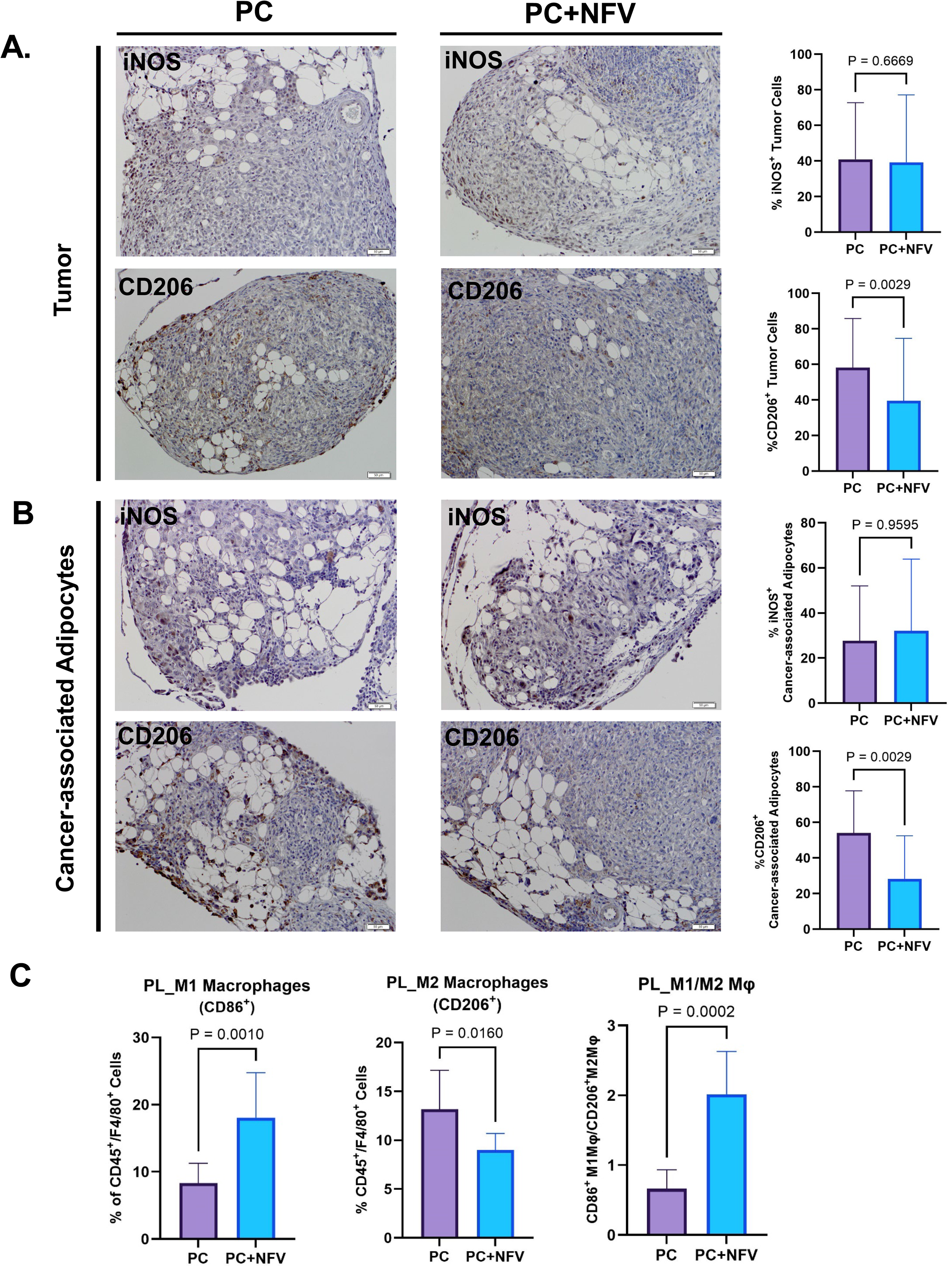
Tumor- and ascites-associated macrophage distribution. Tumor sections were subjected to immunohistochemical staining for iNOS (M1) or CD206 (M2) macrophages. Representative images are shown alongside quantitative graphs with antigen expression in **(A)** tumor and **(B)** intra-tumoral adipose regions using Aperio ImageScope. Pairwise statistical analyses were conducted using Student’s t-test (GraphPad). Scale bar: 50 μm. **(C)** Macrophage profiling of cells collected from ascites fluid was analyzed by multispectral flow cytometry to quantify M1 macrophages (CD86+) and M2 macrophages (CD206+). The complete immune profile is presented in Suppl. Figure 2A.

**Figure 6.**
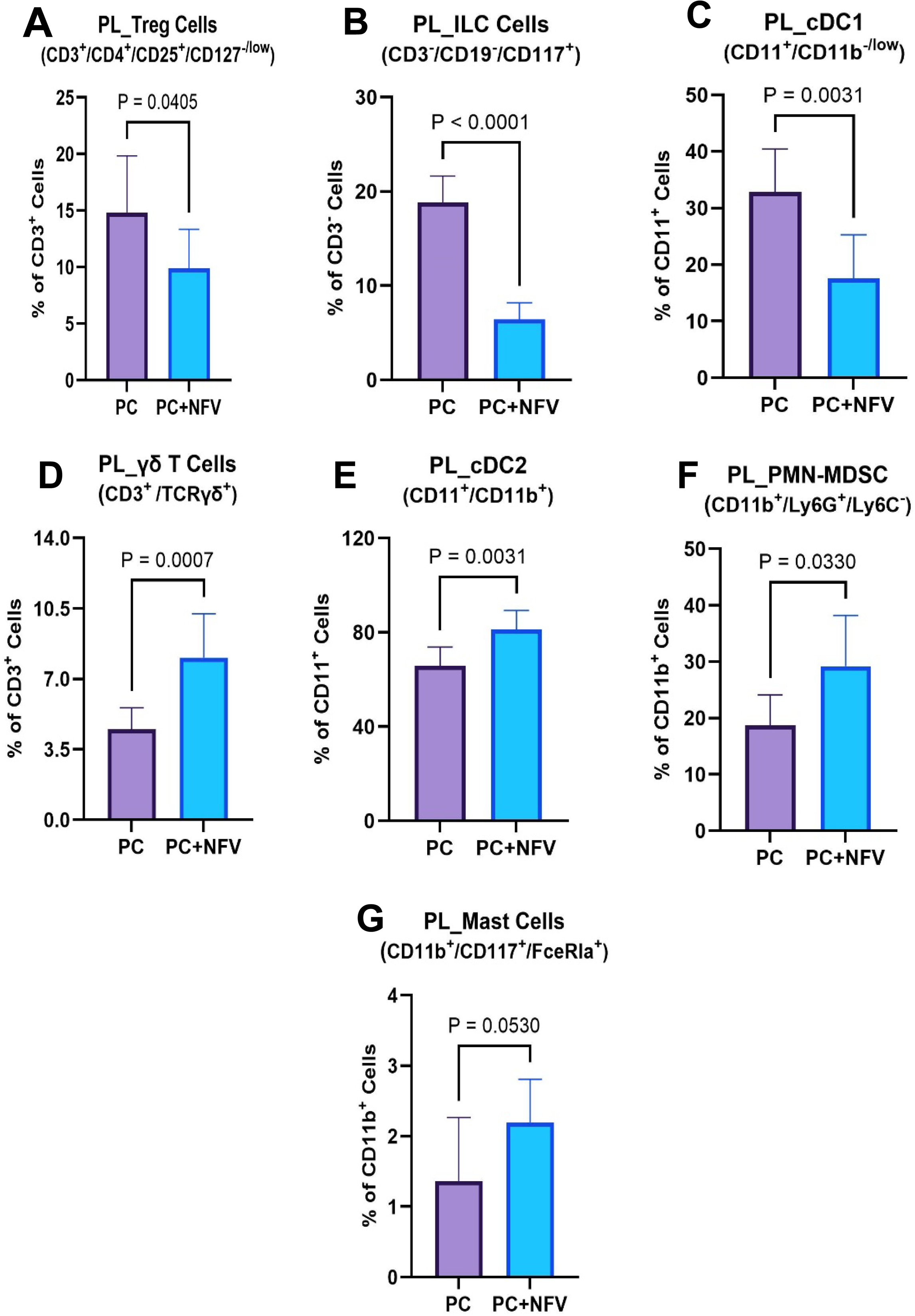
Ascites immune cell landscape. Immune profiling of cells collected from ascites fluid analyzed by multispectral flow cytometry to assess major immune cell subsets. Antibodies are in Suppl. Table 2 and full immune profiles are in Suppl. Fig. 2. **(A)** T-regulatory cells (Tregs), **(B)** innate lymphoid cells (ILCs), **(C)** type 1 dendritic cells (cDC1s), **(D)** γδ-T cells, **(E)** conventional type 2 dendritic cells (cDC2s), **(F)** polymorphonuclear myeloid-derived suppressor cells, and **(G)** mast cells.

## DISCUSSION

Obesity is a recognized noninfectious pandemic (12) that increases OC incidence, enhances metastatic success and reduces survival (13–20). In the U.S., 70% of women are overweight, obese, or extremely obese (34). Meta-analyses, including a study evaluating 2.7 million women (21), show a relationship between obesity and risk of OC incidence in women with tumors of serous, endometrioid, and mucinous histology. Importantly, obesity has an adverse effect on survival of women with OC, implicating a link between host obesity, metastatic success, and response to therapy (12–22). Genomic and clinical data from a cohort of >600 women diagnosed with high grade serous OC (HGSOC) with similar mutational profiles identified a “poor outcome” cohort associated with upregulated obesity- and lipid metabolism-related genes with significantly reduced progression-free and overall survival relative to women with the same mutational profile but lacking this gene cluster (35). These epidemiologic data are consistent with experimental pre-clinical data showing preferential homing of metastasizing ovarian tumors to the omental fat pad (4,5). Data from our lab and others using multiple murine pre-clinical models of diet-induced obesity and mutational obesity have clearly demonstrated that obesity positively correlates with enhanced metastatic success (10,36,37) and is associated with poor response to SOC chemotherapy (11).

Based on data showing enhanced nuclear localization of the transcription factor SREBP1, a master regulator of lipogenesis and lipid transport, in OC tumors from women with high BMI (>30, current study) and in OC tumors from mice fed a high fat diet (10), the current study was designed to test the hypothesis that targeting SREBP1 in combination with SOC chemotherapy would improve outcomes in the high fat diet setting. Pilot studies used two inhibitors reported to block SREBP1 processing: Fatostatin and NFV. While other studies have demonstrated antitumor effects of Fatostatin *in vivo* (38,39), we observed severe dose-limiting toxicity with this compound in the DIO setting and thus discontinued this arm of the trial. As an alternative approach, we repurposed the well-characterized HIV-protease inhibitor NFV (Viracept), which has been demonstrated to also block regulated intramembrane proteolysis catalyzed by site-2 protease (27,28) and importantly displays a well-tolerated safety profile in long-term treatment of human HIV patients over two decades (29). Our results show that NFV in combination with paclitaxel and carboplatin significantly enhanced anti-tumor efficacy and delayed recurrence. This improved therapeutic response likely reflects complementary targeting of both tumor cell proliferation and obesity-associated metabolic pathways, which contribute to tumor progression, immune suppression, and therapeutic response. Nuclear-localized SREBP1 was reduced in both tumor cells and tumor-associated adipocytes in mice treated with PC+NFV, indicating that NFV inhibited SREBP1 processing *in vivo*. Notably, both the number and size of cancer-associated adipocytes were decreased in the combination therapy cohort, providing a potential mechanism for the decreased tumor growth observed in this cohort. These findings highlight the potential of metabolic targeting to augment standard chemotherapy in obesity-associated OC.

Combination therapy with PC+NFV also altered profiles of tumor-associated macrophages in both tissue and ascites relative to mice treated with PC alone, resulting in an increased M1/M2 macrophage ratio primarily due to decreased M2 macrophage polarization. Studies with OC patients show that a high intratumoral M1/M2 macrophage ratio is predictive of improved progression-free survival, platinum-free interval and overall survival (40). Recent studies have shown that alternative (M2) activation of macrophages is triggered in response to interleukin-4 released by T helper cells, that upregulates SREBP1 transcription and target gene expression, with the resulting SREBP1-mediated *de novo* lipogenesis essential for macrophage alternative activation (41). A related study showed that SREBP1 is critical for the survival of M2 tumor-associated macrophages by promoting *de novo* fatty acid synthesis to sustain high energy demands (42). Thus, inhibition of SREBP1 with NFV may directly regulate M2 polarization.

Beyond macrophage remodeling, combination therapy with PC+NFV reprogrammed the ascites immune landscape in obese mice, simultaneously activating multiple anti-tumor pathways while reducing suppressive cell populations. Specifically, combination treatment promoted expansion of M1 macrophages, cDC2s, γδ-T cells, PMN-MDSCs, and mast cells and significantly reduced cDC1s, ILCs, Tregs and M2 macrophages, shifting the ascites microenvironment from an immunologically “cold” to “hot” state. Given that obesity-associated metabolic inflammation impairs dendritic cell function and T-cell priming (43,44), the observed expansion of cDC2s following PC+NFV treatment is consistent with restoration of adaptive immune competence, counteracting immunosuppressive effects typically driven by the obese tumor microenvironment. While cDC1s and cDC2s are not interchangeable (45), reduced cDC1s alongside expanded cDC2s may provide partial functional compensation that supports anti-tumor immunity when CD8⁺ T-cell responses are impaired by obesity.

In parallel, γδ-T cells were selectively expanded. Although γδ-T cells exert potent cytotoxic activity across several tumor models (46–48), their role in ovarian cancer remains poorly defined with reported anti- and pro-tumor functions that are dependent on the tumor microenvironment (49). Notably, anti-tumor γδ-T cell activity thrives in environments with reduced immunosuppressive signals, including low M2 macrophages and Tregs (49), conditions reinforced by PC+NFV-mediated metabolic reprogramming. Despite reductions in total T-cells and no significant changes in CD4^+^ helper or CD8^+^ cytotoxic T cell populations, likely reflecting obesity-driven immunosuppression, the selective expansion of γδ-T cells indicate enhanced innate-like cytotoxic potential, that may compensate for impaired conventional CD8^+^ T-cell responses (50,51). This expansion occurs in the context of reduced Tregs and improved dendritic cell-mediated T-cell priming, reinforcing the shift toward a more cytotoxic and inflammatory microenvironment. It should be noted that T-cell function was not tested in this study. However, Tregs are well-established mediators of immunosuppression in both tumor and ascites microenvironment, with increased accumulation associated with tumor progression and poor survival in ovarian cancer patients (52,53). Mechanistically, Tregs maintain the M2-like macrophage immunosuppression by limiting CD8⁺ T-cell IFNγ production, which normally inhibits SREBP1-driven lipid metabolism (42).

Additional immune changes in ascites after PC+NFV treatment included reduced ILC populations, and increased PMN-MDSCs and mast cells. Reduced ILC populations in ascites may reflect attenuation of tumor-associated inflammatory signaling rather than loss of anti-tumor immunity, although the functional implications of this shift require further investigation (54). Although PMN-MDSCs and mast cells are often considered immunosuppressive (55), the net effect of combination treatment was anti-tumor, as evidenced by the reduction in tumor burden signifying that pro-inflammatory effector populations outweighed potential suppressive influences.

Interestingly, PC + NFV induced distinct compartment-specific immune remodeling. Within the ascites, an antigen-rich and metabolically stressed tumor-associated compartment, combination treatment selectively promoted effector populations and reduced suppressive populations consistent with restoration of local cytotoxic and inflammatory immune function. Systemically, the spleen exhibited reductions in immunosuppressive populations accompanied by increased immune surveillance, representing relief of systemic immunosuppression rather than broad cytotoxic activation. Together, these findings demonstrate that PC+NFV enhances tumor-localized effector activity while maintaining systemic immune balance which may be beneficial in obesity. The induced immune-metabolic reprogramming from PC+NFV treatment establishes a pro-inflammatory microenvironment creating a milieu conducive to tumor clearance, elimination of residual tumor cells and prevention of post-treatment recurrence. Together, these results emphasize the importance of immune-metabolic crosstalk in therapeutic efficacy and support metabolic targeting as a strategy to enhance anti-tumor immunity in obese, immunosuppressed ovarian cancer patients.

A limitation of this study is that NFV has been shown to exert anti-cancer effects by a variety of mechanisms (reviewed in [29]) in addition to inhibition of site-2 protease activity. Thus, while inhibition of SREBP1 processing may have contributed to the enhanced efficacy observed in the combination therapy cohorts, it is likely not the only mechanism. Future studies using ovarian cancer cells and/or murine hosts with targeted deletion of *SREBF1* are needed to delineate the precise contribution of SREBP1. An additional limitation is that the PC+NFV combination therapy was tested only in the obesity-associated ovarian cancer setting. While extensive nuclear SREBP1 staining was not observed in the LFD setting [10] or in tumors from women with low BMI (current study), testing the combination therapy in the LFD setting would help to inform future human clinical trials. To this end, it is interesting to note recent data demonstrating drug synergy between NFV and cisplatin in Pt-resistant ovarian cancer [56], providing additional evidence in support of repurposing NFV for treatment of ovarian cancer.

## Acknowledgement

Some figures are created with BioRender.com.

## Data availability

The data generated in this study are available within the article and its supplementary data files.

