## Supplemental Data for "Metabolic Targeting of SREBP1 Reprograms the Obesity-Driven Ascites Immune Microenvironment and Enhances Chemotherapy Response in Obesity-Associated Ovarian Cancer"

Supplemental Information for: Yang et al.,

**“Combination Therapy Targeting Sterol Regulatory Element Binding Protein 1 (SREBP1)
Improves Response to Standard-of-Care Chemotherapy in Pre-Clinical Models of Ovarian
Cancer and Obesity”**

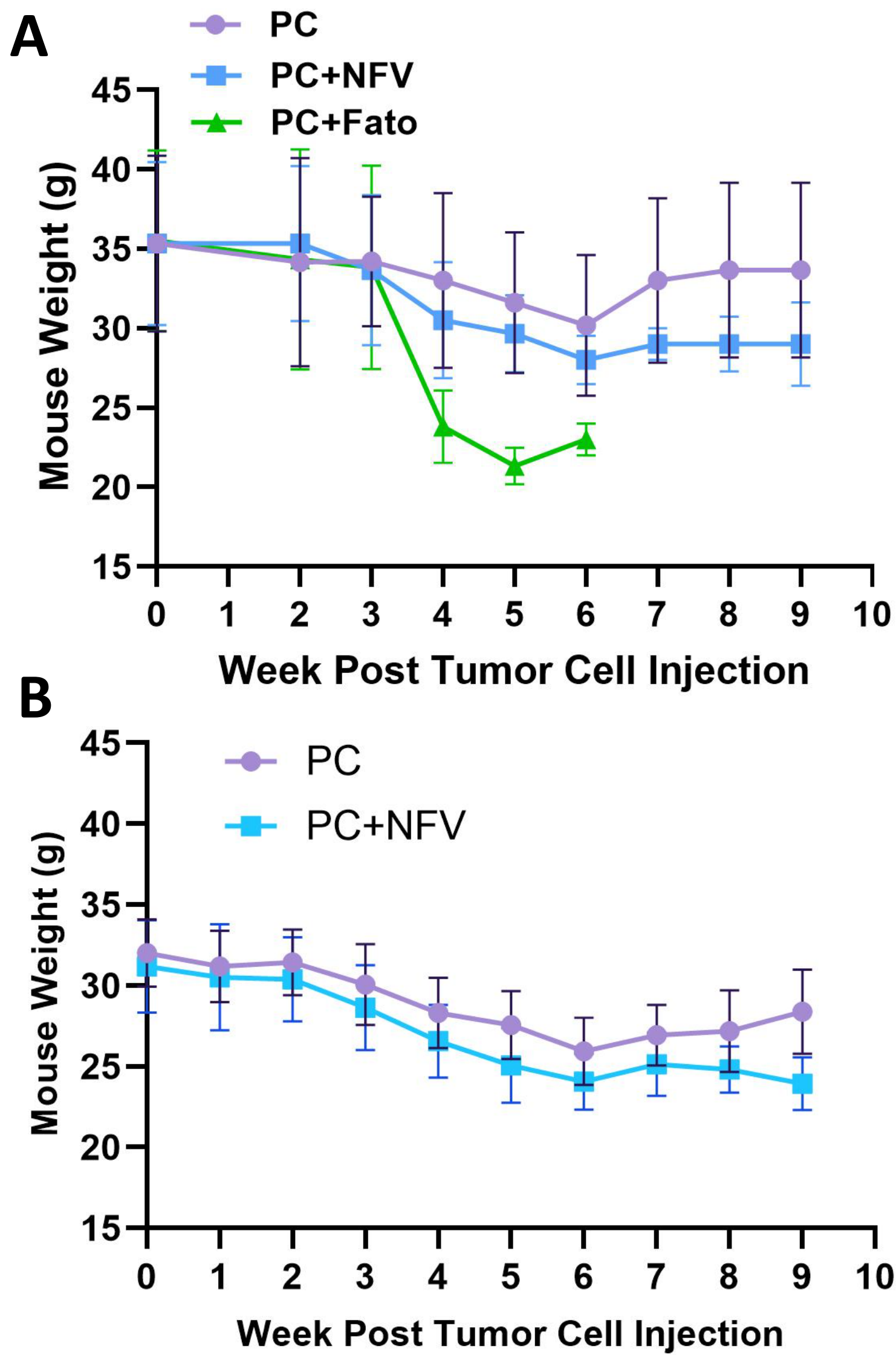

**Suppl Fig. 1.** Mouse body weight post-tumor injection. **(A)** Mice (n=6/cohort) were fed a high fat diet until they reached a weight of 30g prior to tumor cell injection (week 0) as described in Materials and Methods. At week 3, mice were treated with paclitaxel (6mg/kg) + carboplatin (15mg/kg) (designated PC), PC + Nelfinavir (50mg/kg) (PC+NfV) or PC + Fatostatin (30mg/kg) (PC+Fato) twice weekly for 3 weeks. PC+Fato mice experienced significant weight loss compared to the PC and PC+NfV groups, and this arm of the study was discontinued at week 6. Half of the mice in the PC and PC+NfV cohorts were dissected at the end of treatment (week 6) while half were used to evaluate recurrence following treatment cessation (week 9). No significant toxicity was observed with the PC or PC+NfV arms of the study. **(B)** The study was repeated with only PC and PC+NfV cohorts. Mice (n=10/cohort) were fed a high fat diet for 27 weeks prior to tumor cell injection (week 0) as described in Materials and Methods. At week 3, mice were treated at PC as described in (A), or PC+NfV (100mg/kg NfV) for weeks 3-6 followed by NfV alone weeks 7-9. No significant toxicity was observed with the PC or PC+NfV arms of the study.

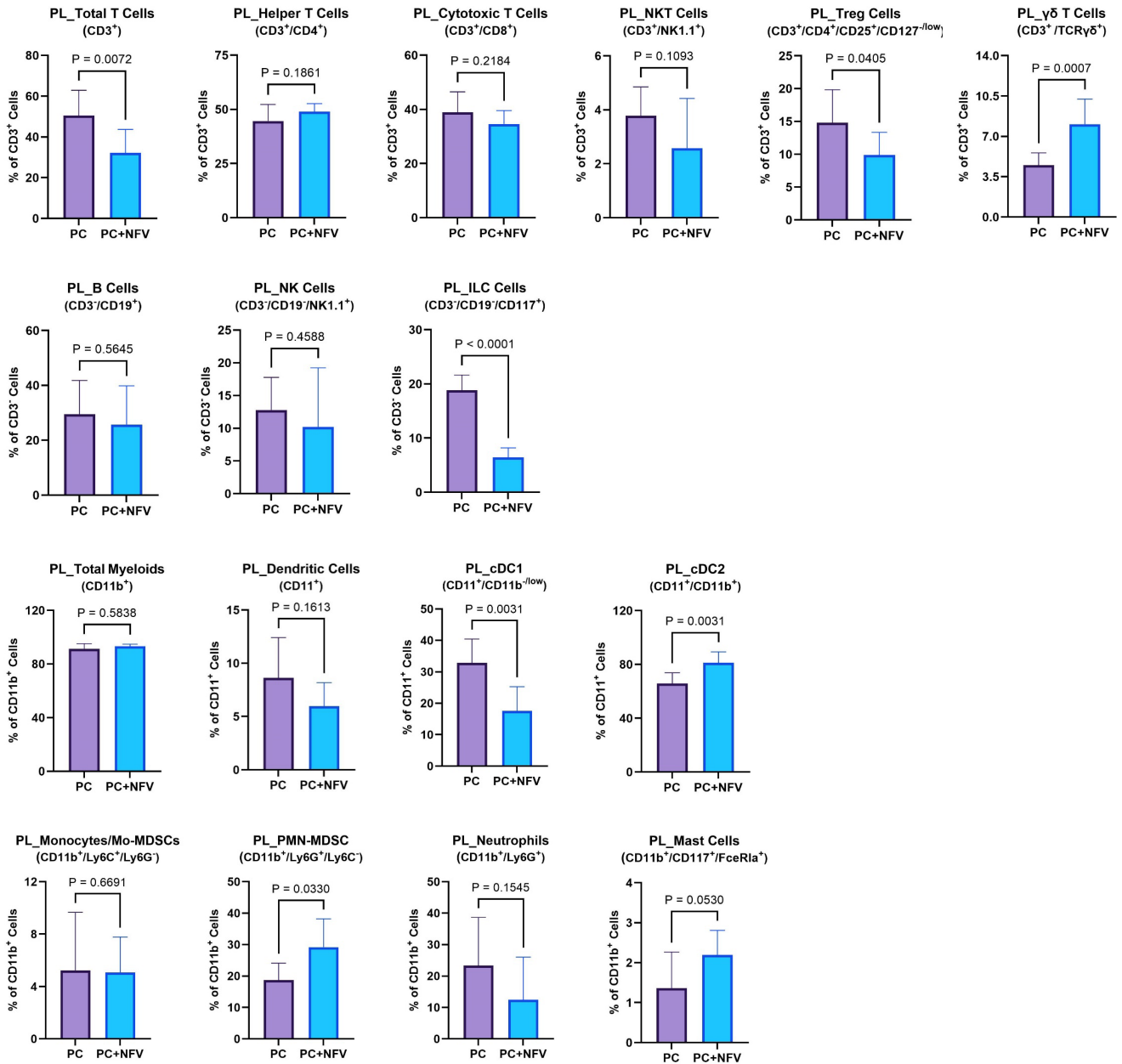

Suppl. Fig. 2A. Flow Cytometry Analysis of Peritoneal Cells.

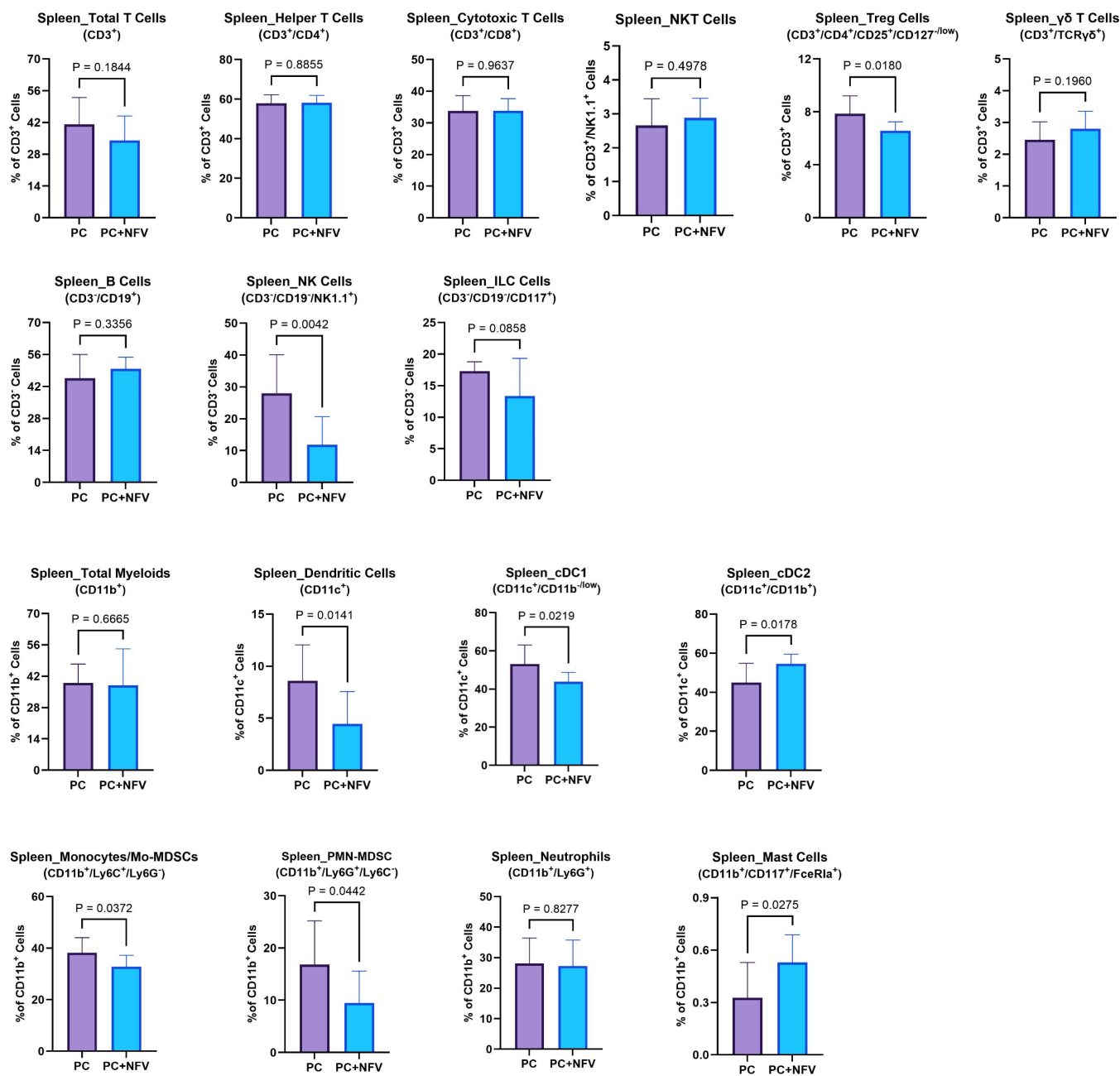

**Suppl. Fig. 2B. Flow Cytometry Analysis of Spleen Cells**

**Suppl Fig. 2. Effect of treatment with the PC + Nelfinavir (PC+NFV) on the peritoneal immune landscape.** Immune profiling of cells collected from peritoneal lavage/ascites **(A)** and the spleen **(B)** were evaluated using the Cyttek Northern Lights™ flow cytometer as described in Materials and Methods using antibodies detailed in Suppl. Table 2.

**(A) Peritoneal immune cells.** The PC+NFV combination therapy group exhibited significantly increased expression of $\gamma\delta$ T cells, cDC2 cells, PMN-MDSCs, and mast cells and significantly decreased total T cells, Tregs, ILC cells, and cDC1 cells compared to the standard of care chemotherapy (PC) cohort. No significant differences were observed upon analyzing expression of helper T cells, cytotoxic T cells, NKT cells, B cells, NK cells, total myeloid cells, dendritic cells, Mo-MDSC or neutrophils.

**(B) Splenic immune cells.** The PC+NFV combination therapy cohort exhibited significantly increased cDC2 cells and mast cells and significantly reduced Tregs, NK cells, dendritic cells, cDC1 cells, Mo-MDSCs, and PMN-MDSCs relative to the chemotherapy-only (PC) cohort. No significant differences were observed for total T cells, helper T cells, cytotoxic T cells, NKT cells, $\gamma\delta$ T cells, B cells, ILCs, total myeloid cells or neutrophils.

| Supplemental Table 1. Patient Information |  |  |  |  |
| --- | --- | --- | --- | --- |
| Patient # | BMI (kg/m<sup>2</sup>) | Age | Diagnosis | Stage |
| 1 | 15.7 | 58 | Carcinoma, NOS | IC |
| 2 | 17.2 | 52 | Serous carcinoma | IIIC |
| 3 | 19.9 | ND | Serous carcinoma | ND |
| 4 | 19.9 | 68 | Papillary serous carcinoma | ND |
| 5 | 20.3 | 79 | Adenocarcinoma, NOS | ND |
| 6 | 20.5 | 56 | Papillary serous adenocarcinoma | IIIC |
| 7 | 21.3 | 65 | Papillary serous carcinoma | ND |
| 8 | 21.4 | 62 | Serous carcinoma | IIIC |
| 9 | 21.7 | 76 | Serous adenocarcinoma | IIIC |
| 10 | 22.4 | 62 | Papillary serous and endometrioid adenocarcinoma | ND |
| 11 | 32.8 | 55 | Serous carcinoma | IIIC |
| 12 | 33.7 | 70 | Serous adenocarcinoma | IIIC |
| 13 | 33.7 | 78 | Papillary serous carcinoma | ND |
| 14 | 35.5 | 53 | Serous carcinoma | ND |
| 15 | 36.3 | 63 | Serous carcinoma | IIIC |
| 16 | 38.4 | 59 | Papillary serous carcinoma | ND |
| 17 | 38.8 | 54 | Papillary serous carcinoma | ND |
| 18 | 42.2 | 52 | Serous adenocarcinoma | ND |
| 19 | 45.9 | 55 | Papillary serous cystadenocarcinoma | ND |
| 20 | 46.9 | 49 | Papillary serous carcinoma | IIIC |

**Supplemental Table 1. Summary of patient data.** Tumor tissues were obtained from patients with normal body mass index (BMI, patients 1-10) or high BMI (patients 11-20). The average BMI of the 'normal BMI' group was 20.03+/-0.66 while the average BMI of the 'high BMI' group was 38.42+/-1.60 (p<0.001). The average age of the 'normal BMI' group was 64.22+/-2.98 while the average age of the 'high BMI' group was 58.80+/-2.87 (p=0.21). (ND=no data; NOS=not otherwise specified)

| Marker | Fluorophore | Company | Catalog # |
| --- | --- | --- | --- |
| CD11c | BB515 | BD Biosciences | 565586 |
| CD4 | BV750 | BD Biosciences | 747344 |
| CD19 | BV711 | BD Biosciences | 563157 |
| FcER1a | BV480 | BD Biosciences | 751769 |
| CD11b | PerCP-Cy5.5 | BioLegend | 101228 |
| CD8 | Spark Blue 550 | BioLegend | 100780 |
| NK-1.1 | BV650 | biolegend | 108736 |
| CD25 | PE | BioLegend | 102008 |
| CD127 | BV605 | BioLegend | 135041 |
| CD117 | PE-Cy7 | BioLegend | 135112 |
| CD16 | efluor 450 | ThermoFisher | 48-0161-82 |
| TCRgd | PerCP-Vio700 | Miltenyi Biotec | 130-104-011 |
| CD3 | AF488 | BioLegend | 100210 |
| Ly6C | BV510 | BioLegend | 128033 |
| LY6G | BV785 | BioLegend | 127645 |
| Tim4 | BV711 | BD Biosciences | 745509 |
| CD31 | BV480 | BD Biosciences | 565629 |
| CD45 | PerCP-Cy5.5 | BioLegend | 103132 |
| CD86 | BV785 | BioLegend | 105043 |
| CD206 | PE | BioLegend | 141706 |
| CD34 | PeCy7 | BioLegend | 128618 |
| F4/80 | efluor 450 | ThermoFisher | 48-4801-82 |

**Supplemental Table 2:** Antibodies used for multiplex flow cytometry
